# Observed Antibody Space: a resource for data mining next generation sequencing of antibody repertoires

**DOI:** 10.1101/316026

**Authors:** Aleksandr Kovaltsuk, Jinwoo Leem, Sebastian Kelm, James Snowden, Charlotte M. Deane, Konrad Krawczyk

## Abstract

Antibodies are immune system proteins that recognize noxious molecules for elimination. Their sequence diversity and binding versatility have made antibodies the primary class of biopharmaceuticals. Recently it has become possible to query their immense natural diversity using next-generation sequencing of immunoglobulin gene repertoires (Ig-seq). However, Ig-seq outputs are currently fragmented across repositories and tend to be presented as raw nucleotide reads, which means nontrivial effort is required to reuse the data for analysis. To address this issue, we have collected Ig-seq outputs from 53 studies, covering more than half a billion antibody sequences across diverse immune states, organisms and individuals. We have sorted, cleaned, annotated, translated and numbered these sequences and make the data available via our Observed Antibody Space (OAS) resource at antibodymap.org. The data within OAS will be regularly updated with newly released Ig-seq datasets. We believe OAS will facilitate data mining of immune repertoires for improved understanding of the immune system and development of better biotherapeutics.

## 1. Introduction

Antibodies (or B-cell receptors) are protein products of B-cells and primary actors of adaptive immunity in jawed vertebrates (1). They are highly malleable molecules that can bind to virtually any antigen. An organism holds a great variety of these molecules increasing the probability that an arbitrary antigen can be recognized by an antibody, initiating an immune response (2). Owing to their binding malleability they are the most prominent class of reagents and biotherapeutics (3, 4). Continued successful exploitation of these molecules relies on our ability to discern the functional diversity of antibody repertoires (5–7).

Next-generation sequencing of immunoglobulin gene repertoires (Ig-seq) has enabled researchers to take snapshots of millions of sequences at a time across individuals, diverse organisms and different immune states (8, 9). The ability to sequence and analyze millions of antibody sequences has the potential to uncover the mechanics of the immune response to any antigen (10, 11) and dysfunctions of the immune system itself (12).

Many previous studies have addressed the issue of antibody diversity, contributing invaluable evidence to understanding the dynamics of human immune systems (13). Numerous analyses have focused on the frequencies of V(D)J gene usages, which can offer insights into creating biased therapeutic antibody libraries (14–16). Another therapeutic application of antibody repertoire analysis is advancing vaccine design by comparative longitudinal studies of pre- and post-antigen challenge experiments (10, 11, 17–22). Such comparative studies have shown that different individuals can converge on the same antibody sequence against a given vaccine (11, 19). Due to sequencing limitations, these analyses have focused on heavy or light chains separately, whereas one ought to study the paired repertoire to obtain deeper insights of antibody diversity (23).

Technical advances in sequencing technology have outpaced storage and analysis pipelines (24, 25). This has meant that the outputs of Ig-seq studies are fragmented across repositories making it difficult to perform large-scale data mining of antibody repertoires (25). Metadata such as isotype, age or subject identifiers are not typically standardized, therefore extraction of specific subsets of antibody repertoires for comparative analyses is challenging. Furthermore, the data are typically deposited as raw nucleotide reads. It requires non-trivial *ad hoc* effort to convert such raw reads to amino acid sequences that ultimately dictate the molecular structure and antigen-recognition. Some of these issues are addressed by services that provide Ig-seq-specific data deposition and analysis pipelines such as Immport (http://immport.org) (26, 27), ImmunoseqAnalyzer (http://clients.adaptivebiotech.com/), IReceptor (http://ireceptor.irmacs.sfu.ca/) or VDJServer (http://vdjserver.org) (28). The IReceptor and the VDJServer are the main resources that fall under the umbrella of the organized effort of the Adaptive Immune Receptor Repertoire (AIRR) Community to provide standardized deposition and analysis pipelines for the Ig-seq outputs (24). These services chiefly focus on facilitating bulk deposition of raw data to perform standardized sequencing analyses. Ultimately, because immunoinformatics is not the chief focus of such services, bulk data download from such websites is limited and converting the raw nucleotide data obtained into a format suitable for analysis still requires non-trivial effort.

To address these issues, we have created the Observed Antibody Space (OAS) resource that allows large-scale data mining of antibody repertoires. We have so far collected the raw outputs of 53 Ig-seq experiments covering over half a billion sequences. We have organized the sequences by metadata such as organism, isotype, B-cell type and source, and the immune status of B-cell donors to facilitate bulk retrieval of specific subsets for comparative analyses. We have converted all of the Ig-seq sequences to amino acids and numbered them using the IMGT scheme. The data is available for querying or bulk download at http://antibodymap.org. We believe that OAS will facilitate data mining antibody repertoires for improved understanding of the dynamics of the immune system and thus better engineering of biotherapeutics.

## 2. Materials and Methods

A list of study accession codes of publically available Ig-seq datasets were obtained via a literature review. The majority of raw reads were downloaded from the European Nucleotide Archive (ENA) (29) and the National Center for Biotechnology Information (NCBI) websites (30). In a small number of cases, another public Ig-seq repository was specified e.g. (14, 31–33). Metadata were manually extracted from the deposited datasets and arranged in a reproducible format.

The downloaded FASTQ files were processed depending on the sequencing platform. Paired raw Illumina reads were assembled with FLASH (34). The assembled antibody sequences were converted to the FASTA format using FASTX-toolkit (35). As raw reads from Roche 454 are not paired, these FASTQ files were directly converted to the FASTA format with the FASTX-toolkit.

The heavy chain sequences were automatically annotated with isotype information unless such data was given in the corresponding publication. Automatic isotype annotation was performed by aligning the constant heavy domain 1 (CH1) of any given antibody sequence against the IMGT isotype reference (36) of the respective species using the Smith-Waterman algorithm (37). We assigned a score of two for a nucleotide match, and a score of minus one for a nucleotide mismatch or a gap. The IMGT isotype references comprised 21 nucleotide-long fragments of the CH1 domain of the antibody isotypes. To ensure a high confidence of correct isotype identification, we employed a conservative threshold of 30 in the Smith-Waterman algorithm scoring function. Sequences whose Smith-Waterman algorithm score was below the threshold for all isotypes were assigned as ‘bulk’. The robustness of this protocol was confirmed on the author-annotated Ig-seq datasets (38–40) where it resulted in 99% accurate annotations. Around 1% of the Ig-seq data had a very short (or missing) of CH1 domain sequence. Such sequences were also assigned as ‘bulk’.

IgBlastn (41) was used to convert the FASTA files of antibody nucleotide sequences to amino acids. The amino acid sequences were then numbered with ANARCI (42) using the IMGT scheme (43). ANARCI does not number a sequence if it does not align to a suitable Hidden Markov Model (44). ANARCI therefore ensures that the antibody sequences do not harbor unusual indels or stop codons in the antibody regions, that the V and J genes align to the respective species amino acid IMGT germlines (36), and that the length of CDR-H3 is not greater than 37 residues in human, mouse, rat, rabbit, alpaca and rhesus antibodies. Due to technical limitations of sequencing platforms, certain reads were missing significant portions of the variable region (e.g. portions of CDR1), sequences that did not have all three CDRs were discarded as incomplete. The V and J genes are identified during the ANARCI numbering step.

Using the protocol above we annotated Ig-seq results of 53 independent studies. In order to streamline updating OAS with new data, we have generated a procedure to automatically identify Ig-seq datasets from raw sequence read archives. We apply our antibody annotation protocol to each raw nucleotide dataset deposited in the NCBI/ENA repositories, if we find more than 10,000 antibody sequences in any given dataset, it is set aside for manual inspection. Manual inspection is still necessary to efficiently assign metadata as these are currently deposited in a non-standardized manner. This procedure allows for automatic identification of new Ig-seq datasets and semi automatically updating of OAS.

## 3. Results

We have so far collected raw sequencing outputs from 53 Ig-seq studies. All raw nucleotide reads were converted into amino acids using IgBlastn (41). The full amino acid sequences were then IMGT numbered using ANARCI (42). As well as providing IMGT and gene annotations, ANARCI acts as a broad-brush filter for antibody sequences that are likely to be erroneous (see Materials and Methods).

Applying the same retrieval, amino acid conversion, gene annotation and numbering protocol to all sequences assures the same point of reference across the 53 heterogeneous Ig-seq datasets (45). This protocol produces the full IMGT-numbered sequences together with gene annotations for each of the datasets.

The numbered amino acid sequences in each dataset are sorted by metadata e.g. individuals, age, vaccination regime, B-cell type and source *etc*. (Figure 1). Deposition of such metadata is currently not standardized and requires *ad hoc* manual curation for each dataset. In an effort to organize the antibody sequences using such metadata, we have grouped the sequences within each dataset into Data Units. Each Data Unit represents a group of sequences within a given dataset with a unique combination of metadata values. The metadata values are summarized in Table 1.

**Figure 1.**
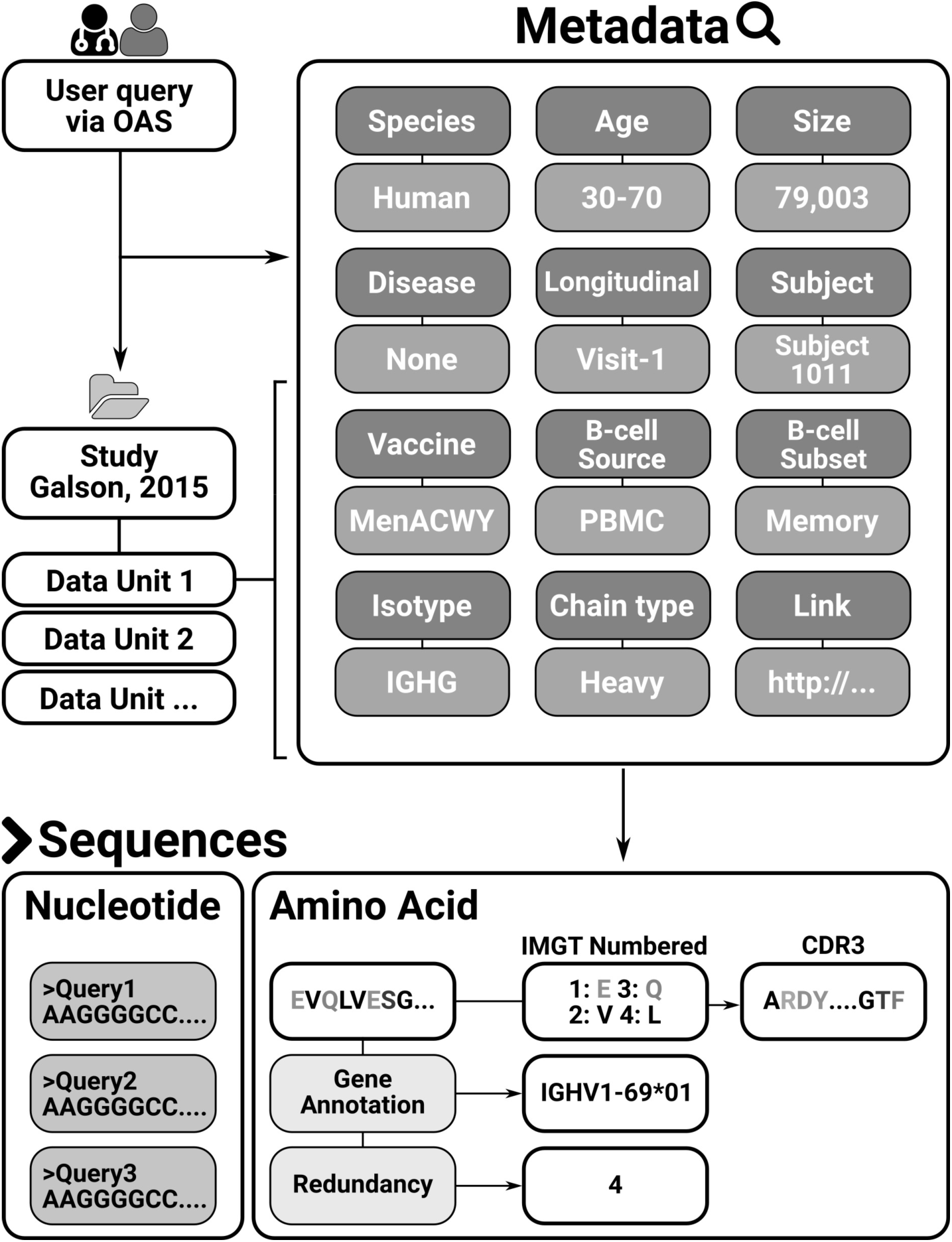
The Observed Antibody Space database. The data from 53 studies is sorted into Data Units. Each Data Unit is a set of antibody sequences that share the same set of meta-data. Each sequence in a Data Unit is further associated with sequence-specific annotations.

**Table 1.** Metadata descriptors of each Data Unit in OAS. Each Data Unit is uniquely identified by the study and a collection of the metadata values.

| Metadata name | Metadata description |
| --- | --- |
| Chain | Heavy/light chain annotation. |
| Isotype | Identified or deposited isotype information. |
| Age | Information on the age of the human B-cell donors. |
| Disease | Indication of whether the donor was sick at the time of B-cell extraction. |
| Vaccine | Indication if the B-cell donor was purposely immunized prior to B-cell extraction. |
| B-cell subset | Indication if a particular B-cell subset was sorted for Ig-seq |
| Species | Organism of the B-cell donor |
| B-cell source | Which organ/tissue the B-cells were extracted from. |
| Subject | Indication of a particular B-cell donor the B-cells were sourced from. |
| Longitudinal | If the study was longitudinal, an indicator of the time point. |
| Size | Number of non-redundant sequences in the dataset. |
| Link | Link to the source publication. |

As of April 29^th^ 2018, 53 Ig-seq studies are included in OAS totaling 608,651,423 sequences (552,824,460 VH and 55,826,963 VL sequences). The majority of these sequences are murine (~50.4%) and human (~47.4%). Twenty-two of the Ig-seq studies interrogate the immune system of diseased individuals, the most common ailment being HIV (13 studies). The database also contains 22 Ig-seq studies of naive antibody gene repertoires (the collection of B-cells from donors who are healthy and not purposefully vaccinated). The main source of B-cells in the OAS database is peripheral blood (~231m of sequences) followed by spleen/splenocytes (~198m) and bone marrow (~124m). The database holds isotype information for each individual heavy sequence and the two most common isotypes are IgM (~312m) and IgG (~139m). For ~65m sequences we were not able to assign isotypes with high confidence. The median total redundant size of the Ig-seq studies in the OSA database is 2,006,196 sequences, while the largest Ig-seq study was that by Greiff et al., (246,449,189 redundant sequences) (14). Detailed statistics on each dataset are given in Table 2. All the data may be bulk downloaded or individual Data Units queried, at http://antibodymap.org.

**Table 2.** Summary of Ig-seq studies that are incorporated into the Observed Antibody Space database.

| Study | Species | Disease | Vaccine | B-cell source | B-cell subset | Total ANARCI<br>parsed<br>sequences |
| --- | --- | --- | --- | --- | --- | --- |
| Banerjee et al.,<br>(46) | Rabbit | None | HIV | PBMC | Unsorted | 4,334,088<br>(2,926,727) |
| Bashford-Rogers et al.,<br>(47) | Human | CLL/None | None | PBMC | Unsorted | 129,013<br>(86,166) |
| Bhiman et al.,<br>(48) | Human | HIV | None | PBMC | Unsorted | 785,751<br>(187,067) |
| Bonsignori et<br>al., (49) | Human | HIV/None | None | PBMC | Memory/<br>Unsorted | 210,377<br>(57,374) |
| Collins et al.,<br>(50) | Mouse | None | None | Splenocytes | Unsorted | 812,439<br>(194,752) |
| Corcoran et al.,<br>(51) | Human/<br>Mouse/<br>Rhesus | None | None | PBMC | Unsorted | 5,307,880<br>(2,840,877) |
| Cui et al., (52) | Mouse | None | NP-CGG/None | Splenocytes | Memory | 5,513,816<br>(935,646) |
| Doria-Rose et<br>al., (13) | Human | HIV | None | PBMC | Unsorted | 2,164,901<br>(549,544) |
| Fisher et<br>al.,(53) | Mouse | None | Plasmodium | Spleen | Unsorted | 175,015<br>(113,594) |
| Galson et al.,<br>(38) | Human | None | Hepatitis-B | PBMC | Unsorted/<br>Plasma cells/<br>HepB-specific | 21,755,739<br>(10,442,291) |
| Galson et al.,<br>(39) | Human | None | Hepatitis-B | PBMC | Unsorted/<br>Plasma cells/<br>HepB-specific | 26,687,394<br>(14,343,236) |
| Galson et al.,<br>(21) | Human | None | Meningitis | PBMC | Naïve/<br>Plasma cells/<br>Memory/<br>Marginal zone | 7,918,197<br>(3,282,907) |
| Galson et al.,<br>(17) | Human | None | Flu | PBMC | Plasma cells | 13,685,210<br>(5,065,786) |
| Greiff et al.,<br>(40) | Mouse | None | NP-CGG | Bone marrow/<br>Spleen | Plasma cells/<br>Plasmablasts | 7,955,739<br>(2,891,649) |
| Greiff et al.,<br>(33) | Mouse | None | NP-CGG | Spleen | ASCs/Plasma<br>cells/<br>Naïve | 788,787<br>(523,716) |
| Greiff et al.,<br>(14) | Mouse | None | OVA/Hepatitis-B/<br>NP-HEL/None | Spleen/<br>Bone marrow | Plasma cells/<br>Pre-B-cells/<br>Naïve | 246,449,189<br>(129,417,638) |
| Gupta et al.,<br>(54) | Human | None | Flu/Hepatitis-A/<br>Hepatitis-B | PBMC | Unsorted | 25,134,322<br>(9,966,175) |
| Halliley et al.,<br>(55) | Human | None | Flu/Tetanus | Bone marrow | Plasma cells | 2,348,164<br>(1,208,616) |
| Huang et al.,<br>(56) | Human | HIV | None | PBMC | Memory | 11,693,783<br>(5,701,433) |
| Jiang et al., (57) | Human | None | Flu | PBMC | Naïve/<br>Plasmablasts | 3,199,271<br>(1,809,306) |
| Joyce et al.,<br>(58) | Human | None | None | PBMC | Unsorted | 2,747,688<br>(1,463,421) |
| Khan et al., (59) | Mouse | None | OVA | Spleen | Unsorted | 24,175,033<br>(7,113,411) |
| Levin et al., (60) | Human | Allergy | None | PBMC/<br>Nasal biopsy | Unsorted | 528,173<br>(370,465) |
| Levin et al., (61) | Human | Allergy | None | PBMC/<br>Bone marrow | Unsorted | 29,643,305<br>(9,557,586) |
| Li et al., (62) | Camel | None | None | PBMC | Unsorted | 1,152,359<br>(1,127,651) |
| Liao et al., (63) | Human | HIV | None | PBMC | Unsorted | 1,420,314<br>(619,492) |
| Lindner et al.,<br>(64) | Mouse | None | E.Coli/<br>Clostridia/<br>Lactobacillus | Biopsy of small<br>intestine | Unsorted | 1,686,350<br>(544,061) |
| Meng et al.,<br>(65) | Human | CMV/<br>EBV/None | None | PBMC/Lung/<br>Spleen/Bone<br>marrow/Colon/<br>Jejunum/Lymph<br>node/Ileum | Unsorted | 45,576,606<br>(21,738,501) |
| Menzel et al.,<br>(66) | Mouse | None | NP-CGG | Spleen/Bone<br>marrow | ASCs | 14,355,151<br>(6,058,480) |
| Mroczek et al.,<br>(67) | Human | None | None | PBMC | Immature/<br>Transitional/<br>Mature/<br>Plasmacytes/<br>Memory | 104,154<br>(85,525) |
| Ota et al., (68) | Mouse | None | None | Spleen/Lymph | Unsorted | 21,505<br>(9,619) |
| Palanichamy et<br>al., (69) | Human | MS | None | CSF/PBMC | Unsorted | 776,895<br>(292,801) |
| Parameswaran<br>et al., (11) | Human | Dengue/None<br>/ Non-dengue<br>febrile illness/ | None | PBMC | Unsorted | 26,584<br>(23,606) |
| Prohaska et al.,<br>(70) | Mouse | None | None | Spleen/<br>Peritoneum | B-1/ B-2/<br>Marginal Zone/<br>Follicular | 336,723<br>(198,983) |
| Rettig et al.,<br>(32) | Mouse | None | None | Spleen/<br>Splenocytes | Unsorted | 41,908<br>(24,908) |
| Rubelt et al.,<br>(71) | Human | None | None | PBMC | Naïve/Memory | 2,320,947<br>(1,719,507) |
| Schanz et al.,<br>(31) | Human | HIV/None | None | PBMC | Unsorted | 12,734,958<br>(5,412,549) |
| Zhu et al.,<br>(72) | Human | HIV | None | PBMC | Unsorted | 1,962,643<br>(532,350) |
| Stern et al.,<br>(73) | Human | MS | None | Cervical lymph<br>node/<br>White matter<br>lesion/ Pia<br>mater/ Choroid<br>plexus/ Cortex/<br>Spleen | Unsorted | 8,550,247<br>(3,321,530) |
| Sundling et al.,<br>(74) | Rhesus | None | HIV | PBMC | Unsorted | 40,960<br>(26,298) |
| Tipton et al.,<br>(75) | Human | SLE/None | Flu/Tetanus | PBMC | Unsorted | 28,204,742<br>(13,301,396) |
| Turchaninova<br>et al., (76) | Human | None | None | PBMC | Memory/Plasma<br>cells/Naïve | 183,949<br>(176,441) |
| Vander Heiden<br>et al., (77) | Human | MG/None | None | PBMC | Memory/Naïve/<br>Unsorted | 13,939,166<br>(5,170,299) |
| VanDuijn et al.,<br>(78) | Rat | None | DNP/HuD | Splenocytes | Unsorted | 6,359,396<br>(4,234,597) |
| Vergani et al., (79) | Human | None | None | PBMC | Unsorted | 14,161,949<br>(5,987,086) |
| Wasemann et al., (80) | Mouse | None | NP-CGG | Lamina propria/<br>Bone marrow/<br>Spleen | Unsorted | 146,370<br>(40,132) |
| Wu et al., (81) | Human | HIV | None | PBMC | Unsorted | 394,144<br>(198,468) |
| Wu et al., (82) | Human | HIV | None | PBMC | Unsorted | 5,545,910<br>(1,370,109) |
| Wu et al., (83) | Human | Allergy/ None | None | PBMC/<br>Nasal biopsy | Unsorted | 35,034<br>(23,923) |
| Zhou et al., (22) | Human | HIV | None | PBMC | Unsorted | 1,541,645<br>(458,227) |
| Zhou et al., (84) | Human | HIV | None | PBMC | Unsorted | 722,112<br>(291,670) |
| Zhu et al., (85) | Human | HIV | None | PBMC | Unsorted | 874,930<br>(174,435) |
| Zhu et al., (86) | Human | HIV | None | PBMC | Unsorted | 1,290,499<br>(699,828) |

The datasets are organized into studies related to a given Ig-seq experiment. Each study in the OAS database is subdivided into Data Units. Each Data Unit is a collection of IMGT-numbered amino acid sequences uniquely identified by the metadata descriptors given in Table 1, five of which (species, disease, vaccine, B-cell source and B-cell type) are given in this Table. The ‘total ANARCI parsed sequences’ field indicates the total number of redundant sequences in our database, with the non-redundant numbers in parentheses. Abbreviations: PBMC, peripheral blood mononuclear cell; CLL, chronic lymphocytic leukemia; NP-CGG, chicken gamma globulin; ASC, antibody secreting cell; OVA, ovalbumin; NP-HEL, hen egg lysozyme; CSF, cerebrospinal fluid; MS, multiple sclerosis; SLE, systemic lupus erythematosus; MG, myasthenia gravis; DNP, dinitrophenyl; HuD, paraneoplastic encephalomyelitis antigen.

## 4. Discussion

Here, we describe the Observed Antibody Space (OAS) database, a unified repository to facilitate large-scale data mining of antibody repertoires in both their amino acid and nucleotide forms. Absence of well established repositories in Ig-seq deposition space required us to perform a combination of literature search and manual curation of the datasets in order to organize the data into OAS. The current lack of widely-adopted deposition standards hampers automatic updating of OAS, as datasets where we find large number of antibodies still require manual curation to perform metadata annotation correctly. Hopefully, efforts such as that by the AIRR community will result in standardization of Ig-seq outputs and will further streamline deposition procedures facilitating large-scale data mining of antibody repertoires (24). Devising unified antibody repertoire repositories is challenging due to both the size of the datasets as well as the diverse data descriptors and analytical pipelines desired by bioinformaticians, wetlab scientists and clinicians (33).

OAS is the first organized collection of a large body of Ig-seq outputs. In order to allow comparative bioinformatics analyses across different subsets of antibody repertoires, we have annotated the datasets by commonly used metadata descriptions such as organism, isotype, B-cell type and source, and the immune state of B-cell donors. To facilitate research about particular antibody sequences or regions, we make full IMGT-numbered high-quality amino acid sequences available together with gene annotations, as well as raw nucleotide data.

This data should aid in-depth comparative analyses across different studies to discern the commonalities observed between independent samples as well as directing Ig-seq experiments on not yet interrogated antibody repertoires. Revealing shared preferences can be invaluable in identifying the portions of the theoretically allowed antibody space that are strategically used to start immune responses (6). Furthermore, such comparative studies can offer a way of deconvoluting the various degrees of freedom of immune repertoires such as differences between diversity of isotypes (67) or organisms (87). Charting the differences between repertoires of human/mouse is of particular interest for engineering better humanized biotherapeutics (88).

Beyond identifying broad commonalities across repertoires, data mining Ig-seq outputs provides novel avenues for designing better antibody-based therapeutics. The plethora of currently available Ig-seq data offers a glimpse at a set of sequences that should be able to fold and function in an organism. Aligning therapeutic candidates to sequences in Ig-seq repertoires can reveal mutational choices that might be naturally acceptable hence providing shortcuts for antibody engineering such as humanization (89). Furthermore, contrasting the naturally observed antibodies with therapeutic ones can offer insight as to the naturally favored biophysical properties of these molecules (4). All such future applications rely on the availability of well-structured datasets that can offer a unified point of reference for bioinformatics analyses. We hope that OAS will aid data mining antibody repertoires, help identify strategic preferences of our immune systems and will ultimately improve how we engineer antibodies into better therapeutics.

## Acknowledgment

We would like to thank all members of Oxford Protein Informatics Group for testing our OAS resource. In particular, we are grateful to Garret M. Morris and Matthew Raybould for their comments, which significantly improved the quality of our work.

## Funding

This work was supported by funding from Biotechnology and Biological Sciences Research Council (BBSRC) [BB/M011224/1] and UCB Pharma Ltd awarded to AK.

